# The third-generation tetracycline KBP-7072 exploits and reveals a new potential of the primary tetracycline binding pocket

**DOI:** 10.1101/508218

**Authors:** Tatsuya Kaminishi, Andreas Schedlbauer, Borja Ochoa-Lizarralde, Elisa de Astigarraga, Retina Çapuni, Fred Yang, Vincent Benn, Qingmei Liu, Xiaojuan Tan, Min Zhang, Sean R. Connell, Paola Fucini

**Affiliations:** Molecular Recognition and Host-Pathogen Interactions, CIC bioGUNE, Parque Científico y Tecnológico de Bizkaia, 48160 Derio, Bizkaia, Spain; KBP Biosciences USA Inc., Princeton, NJ, USA; KBP Biosciences Co., Ltd., Jinan, Shandong, China; IKERBASQUE, Basque Foundation for Science, 48013 Bilbao, Spain

**Author notes:** These authors contributed equally to the work (Tatsuya Kaminishi, Andreas Schedlbauer).

**Keywords:** KBP-7072, antibiotics, tetracycline, ribosome, X-ray crystallography, protein synthesis

## Abstract

Antibiotic resistance is a growing threat to human health requiring the discovery or development of new anti-infectives. As such, KBP-7072 is a novel tetracycline derivative that exhibits broad-spectrum activity against Gram-positive and -negative bacterial strains. To determine the mechanism of action of KBP-7072 and understand how its unique C9 extension can be used to combat the growing problem of antibiotic resistance we determined the structure of KBP-7072 bound to the bacterial 30S ribosomal subunit, the inhibitory target of typical tetracyclines. We show that KBP-7072 binds to the primary tetracycline binding site on the 30S ribosomal subunit consistent with it acting as a protein synthesis inhibitor that blocks A-site occupation. Moreover, the unique chemical nature of KBP-7072´s C9 extension leads to a distinctive interaction pattern with the 30S subunit that distinguishes KBP-7072 from the third-generation tetracycline, Tigecycline, and thus expands the interaction potential of the primary tetracycline binding pocket.

## INTRODUCTION

The World Health Organization has declared antimicrobial resistance a “serious threat [that] is no longer a prediction for the future, it is happening right now in every region of the world and has the potential to affect anyone, of any age, in any country” (1). Overcoming the threat posed by increasing antimicrobial resistance hinges on our ability to introduce into the clinical setting novel and/or more effective antimicrobial agents by implementing new strategies for their discovery and development. Derivatization of known classes of antibiotics offers a solid basis for this search as the parental molecules are well characterized in terms of their pharmacological properties, mechanism of action and resistance mechanisms. The tetracycline class of antibiotics, are one such example, as they have, since their discovery in 1948-1953, proven to be safe and effective against a broad spectrum of Gram positive and negative bacteria and, accordingly, have been in continuous clinical use for more than sixty years (2, 3). Moreover, the mode of action of typical tetracyclines is well known and they function by binding to the bacterial ribosome, acting as protein synthesis inhibitors.

Today wide spread bacterial resistance is compromising the effectiveness of many tetracyclines used clinically and has spurred advances in their chemical synthesis paving the way for the discovery of more potent derivatives (2-4). One of the most effective modifications discovered so far involves alterations at the C9 position of the naphthacene core. This is, for example, the case of Tigecycline (TIG; **Figure 1**), the first semisynthetic third generation tetracycline derivative approved by the FDA and in use in clinical settings since 2005 (5), as well as the case of more recently discovered tetracycline derivatives, i.e. TP-271(6) and TP-434 (7), currently in clinical trials. TIG, TP-271 and TP-434 are all characterized by a unique extension at the C9 position and in comparison, to the native tetracycline molecule, they present a higher antimicrobial efficiency of 10-100 times (6, 7). Additional biochemical and biophysical studies aimed at comparing Tigecycline and other 3rd generation Tetracycline derivatives, including Fluorocycline, Azatetracycline, Omadacycline and Pentacycline, indicate that the derivative’s efficiency can be rationalized in terms of its mode of binding to the 30S ribosomal subunit (8, 9). In this regard, structural studies, conducted so far only on tetracycline (10, 11) and Tigecycline (8, 9, 12), have revealed that although Tigecycline interacts with the same pocket on the ribosome as Tetracycline, the sidechain extension at the C9 position of TIG establishes additional contacts, via the glycyl nitrogen and the secondary amine present in the 9-*tert*-butylglycylamide moiety, with the ribosomal nucleotide C1054. These additional interactions can rationalize the increased affinity observed for TIG in comparison to TET, as well as the ability of TIG to overcome resistance mechanisms mediated by the ribosomal protection protein Tet(M) (9). Namely, the *tert*-butylglycylamide substituent of TIG sterically hinders Tet(M) from correctly accessing the tetracycline binding site and prevents TetM from promoting the release of the bound antibiotic (8, 9, 13).

**Figure 1:**
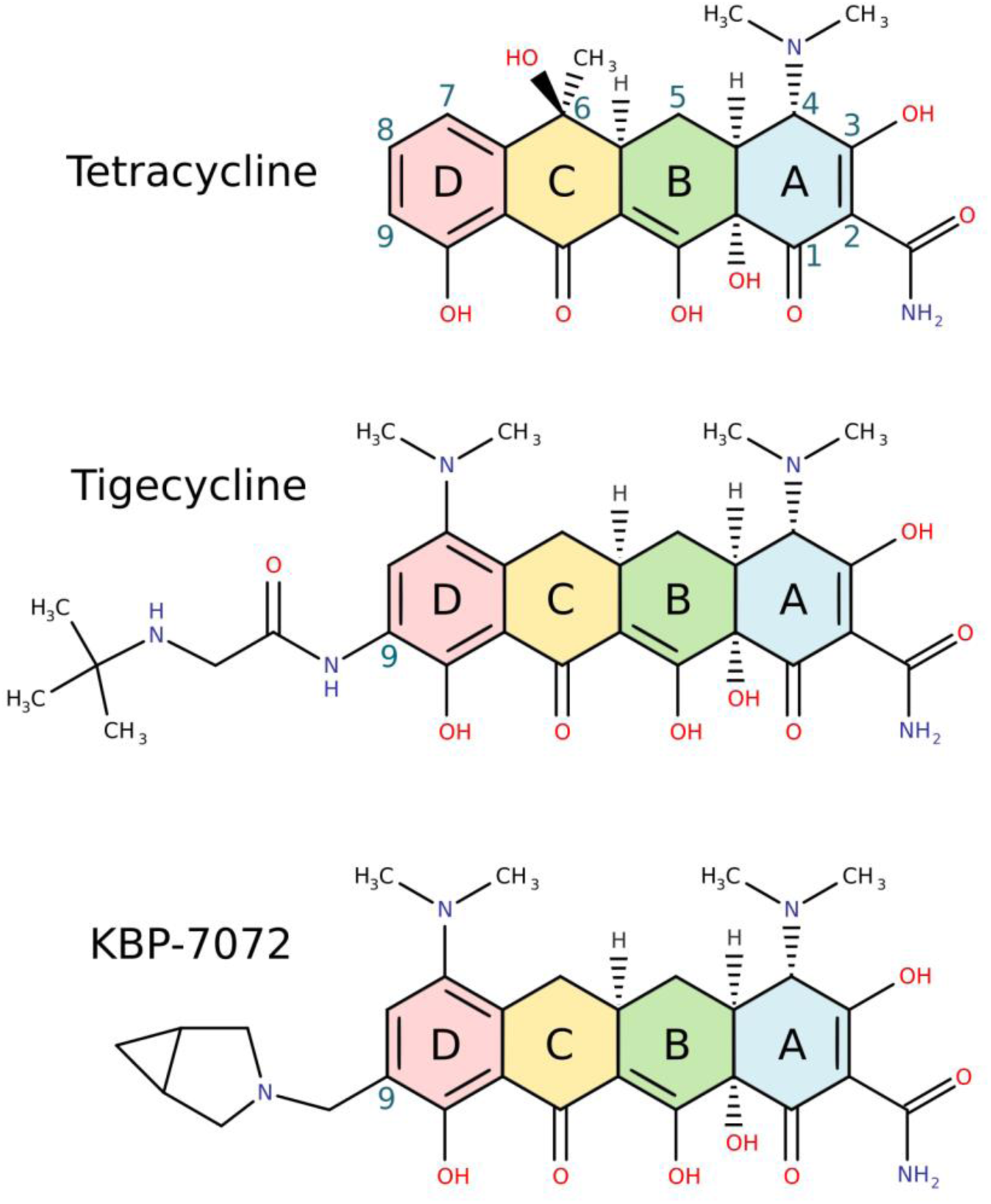
The chemical structures of tetracycline, tigecycline and KBP-7072 are drawn schematically with their common backbone structures (rings A–D) colored distinctly. Carbon atom assignments for the 4-ring backbone (naphthacene core) are indicated on tetracycline. Tigecycline and KBP-7072 differ in the C9 substituent, a *tert*-butylglycylamide sidechain and a 3-methyl-3-azabicyclo[3.1.0]hexane sidechain, respectively.

To further our understanding of the molecular foundations dictating the interaction of the C9 extension with the ribosome, in this study, we characterize the binding mode of a novel and well-tolerated third-generation tetracycline derivative, KBP-7072 (**Figure 1**) (14-16), on the bacterial 30S ribosomal subunit. As an antibiotic KBP-7072 demonstrates potent *in vitro* activity against *Staphylococcus aureus* and *Streptococcus pneumoniae* strains resistant to methicillin, penicillin, and tetracycline (17) and shows a linear dose response exposure, in humans, in single dose or multiple dose studies (15, 16). Using X-ray crystallography, we have determined the structure of KBP-7072 in complex with the *Thermus thermophilus* 30S ribosomal subunit revealing that KBP-7072 binds within the primary tetracycline binding site where it would sterically block aminoacyl-tRNA accommodation as typical members of the tetracycline class of antibiotics. The resulting structure provides a description of the interactions that the unique 3-methyl-3-azabicyclo[3.1.0]hexane sidechain of KBP-7072 makes with the 30S ribosomal subunit. Interestingly, these interactions are different from the ones observed for the functional C9 extension of TIG, providing a structural basis to further investigate possible differences in the activity of TIG and KBP-7072 and a new structural framework for the preparation of even more potent derivatives.

## MATERIALS AND METHODS

### Ribosome preparation and crystallization

Starting inoculum for *T. thermophilus* HB8 (DSM-579) cell cultures were purchased from the Deutsche Sammlung von Mikroorganismen und Zellkulturen GmbH (DSMZ, Germany). The 30S ribosomal subunits were purified from *T. thermophilus* HB8 and crystallized using established methods (8, 18). Crystals were soaked for 12 to 24 h with 17 μM KBP-7072, provided by KBP BioSciences Co., Ltd., and flash-frozen in liquid nitrogen in the presence of the cryoprotectant 2-methyl-2,4-pentanediol.

### Data Collection and Processing

X-ray diffraction data were collected at three synchrotron radiation facilities, namely the XALOC beam line at ALBA (Spain), the i04-1 beam line at Diamond Light Source (UK), and the id29 and id30b beam lines at the European Synchrotron Radiation Facility (France). The diffraction data were processed with the XDS (19) and CCP4 (20) program packages. The native structure of the 30S subunit (PDB code 2ZM6) was refined against the structure factor amplitudes of the 30S·KBP-7072 complex using the PHENIX program package (21). KBP-7072 was modelled into residual electron density using COOT (22) and the resulting 30S·KBP-7072 model was further refined to convergence. For the calculation of the free R factor, 5% of the data were omitted throughout refinement. Figures containing structures are illustrated with MarvinSketch (ChemAxon; https://chemaxon.com/)), inkscape (https://inkscape.org/en/) and PyMOL (http://pymol.sourceforge.net). rRNA residues are numbered according to *the Escherichia coli* scheme, and helices are indicated using the standard nomenclature (23) throughout the article.

## RESULTS

### KBP-7072 binds the primary tetracycline binding site

To elucidate the interaction of KBP-7072 with the ribosome, we determined the crystal structure of the antibiotic bound to the *T. thermophilus* 30S ribosomal subunit, a model system previously used to investigate the binding mode of a variety of antibiotics, among which are Tetracycline and Tigecycline (8, 10, 18, 23, 24). As seen in **Figure 2**, in the initial unbiased Fo-Fc difference Fourier map, positive electron density (green mesh, **Figure 2B**) is readily observed in the primary tetracycline binding site, a pocket formed on the 30S ribosomal subunit by helix 34 (blue) and the loop of helix 31 (red; **Figure 2A**). Importantly the initial unbiased Fo-Fc difference Fourier map fully accommodates the complete KBP-7072 molecule (**Figure 2B)** including the 4-ring backbone that is common to all tetracycline derivatives, as well as the 3-methyl-3-azabicyclo[3.1.0]hexane sidechain that is unique to KBP-7072. Although KBP-7072 was soaked into the 30S subunit crystal using relatively high concentrations (17-44 μM) KBP-7072 was observed to be bound at only the primary tetracycline binding site. This suggests that KBP-7072, like Tigecycline, shows higher binding specificity to the 30S subunit than the parent compound, Tetracycline, which has been observed at several secondary binding sites (10-12, 25). As this initial unbiased Fo-Fc difference Fourier map unambiguously established the presence of KBP-7072 in the ribosomal complex, the KBP-7072 structure was added to the 30S model and subsequent refinement at the resolution of 3.1 Å led to a final structure with crystallographic Rwork/Rfree values of 19.3%/23.4% (**Table 1**). As seen in **Figure 2C** the final 2F_o_-F_c_ electron density map (blue mesh) is of high quality and clearly delineates the shape of the KBP-7072 model including its unique side chain. The completeness of the density attributable to the KBP-7072 molecule, even readily observed in the initial Fo-Fc map (**Figure 2B**), indicates that the entire KBP-7072 molecule is well ordered and structured in its bound form in the 30S·KBP-7072 complex.

**Table 1:**
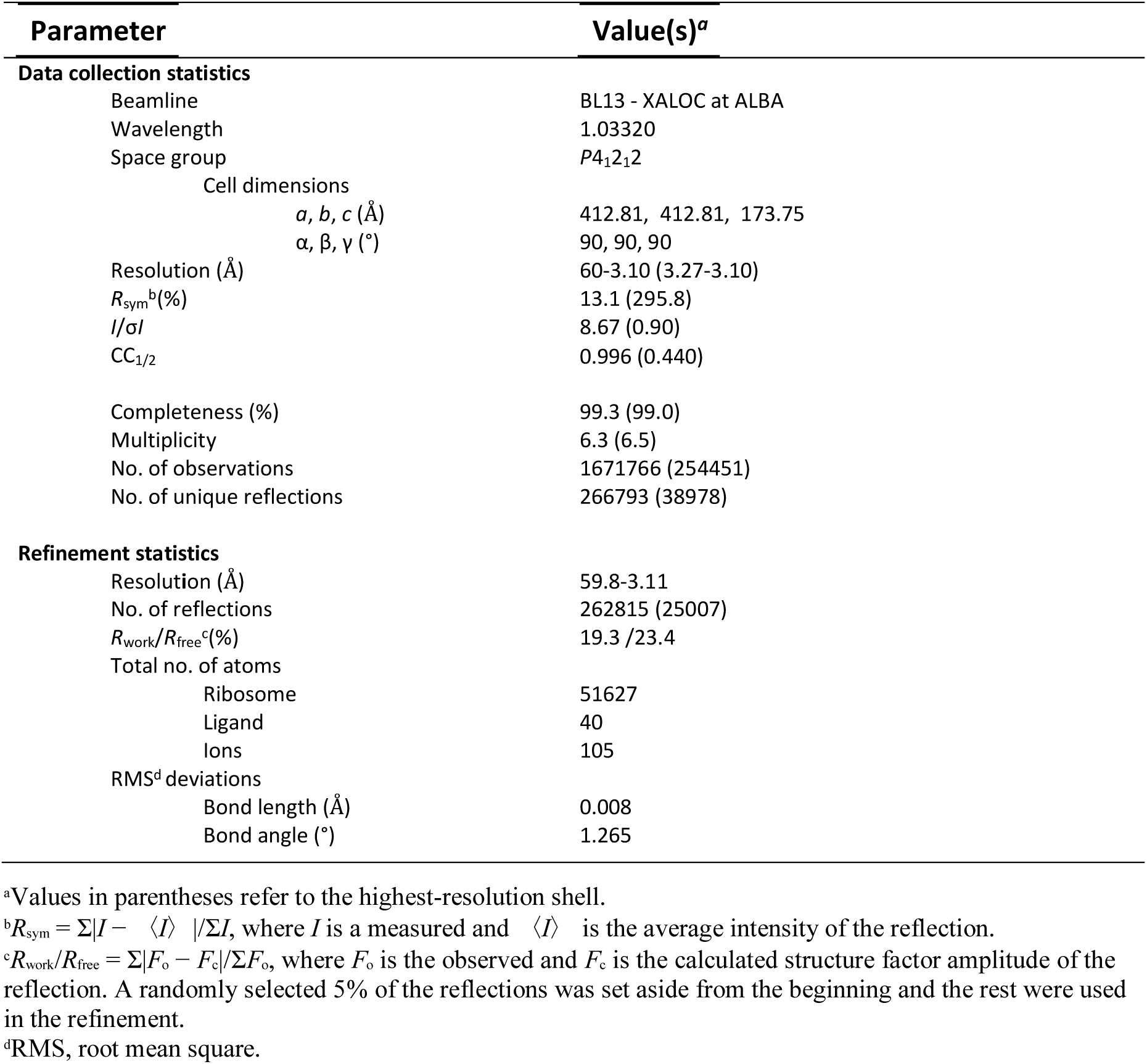
Data collection and refinement statistics

| Parameter | Value(s)<sup>a</sup> |
| --- | --- |
| **Data collection statistics** |  |
| Beamline | BL13 - XALOC at ALBA |
| Wavelength | 1.03320 |
| Space group | *P*4<sub>1</sub>2<sub>1</sub>2 |
| Cell dimensions |  |
| *a*, *b*, *c* (Å) | 412.81, 412.81, 173.75 |
| $\alpha$, $\beta$, $\gamma$ (°) | 90, 90, 90 |
| Resolution (Å) | 60-3.10 (3.27-3.10) |
| *R*<sub>sym</sub><sup>b</sup>(%) | 13.1 (295.8) |
| *I*/σ*I* | 8.67 (0.90) |
| CC<sub>1/2</sub> | 0.996 (0.440) |
| Completeness (%) | 99.3 (99.0) |
| Multiplicity | 6.3 (6.5) |
| No. of observations | 1671766 (254451) |
| No. of unique reflections | 266793 (38978) |
| **Refinement statistics** |  |
| Resolution (Å) | 59.8-3.11 |
| No. of reflections | 262815 (25007) |
| *R*<sub>work</sub>/*R*<sub>free</sub><sup>c</sup>(%) | 19.3 /23.4 |
| Total no. of atoms |  |
| Ribosome | 51627 |
| Ligand | 40 |
| Ions | 105 |
| RMS<sup>d</sup> deviations |  |
| Bond length (Å) | 0.008 |
| Bond angle (°) | 1.265 |

<sup>a</sup>Values in parentheses refer to the highest-resolution shell.
<sup>b</sup> $R_{\text{sym}} = \sum |I - \langle I \rangle| / \sum I$ , where *I* is a measured and $\langle I \rangle$ is the average intensity of the reflection.
<sup>c</sup> $R_{\text{work}}/R_{\text{free}} = \sum |F_o - F_c| / \sum F_o$ , where *F*<sub>o</sub> is the observed and *F*<sub>c</sub> is the calculated structure factor amplitude of the reflection. A randomly selected 5% of the reflections was set aside from the beginning and the rest were used in the refinement.
<sup>d</sup>RMS, root mean square.

**Figure 2:**
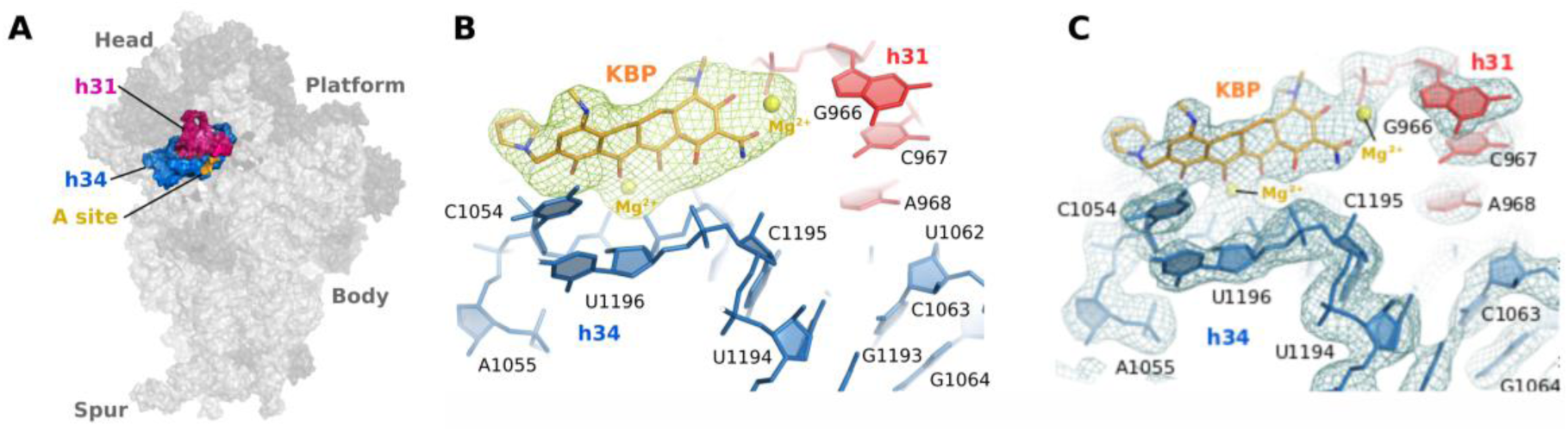
The position of the primary tetracycline binding site is shown on the structure of the 30S ribosomal subunit (**A**) with 16S rRNA elements forming the binding pocket, namely helix 31(h31) and helix 34 (h34) colored red and blue, respectively. This site overlaps with the binding site for aminoacyl-tRNA (A-tRNA; important residues for A-tRNA binding are orange). In panel **B**, the initial Fo-Fc difference Fourier map (colored in green and contoured at 3s) from the 30S-KBP-7072 structure indicates the presence of an additional mass within the primary tetracycline binding site (h31, red; h34, blue). This initial electron density map readily accommodates KBP-7072 (orange sticks) including, importantly, the 3-methyl-3-azabicyclo[3.1.0]hexane sidechain moiety that differentiates KBP-7072 from other tetracycline derivatives. In panel **C** the final 2F_o_-F_c_ electron density map (blue mesh) is shown contoured at 1 sigma and sharpened using a B-factor of -80. KBP-7072 and 16S rRNA residues corresponding to h31 and h34 are colored as in panel **B**.

### Interaction of KBP-7072 with the primary Tetracycline binding site

The atomic model of the 30S·KBP-7072 complex allows for the characterization of the interaction of the KBP-7072 molecule with the ribosomal binding pocket (**Figure 3**) and the comparison of the identified interactions with those reported for Tetracycline and Tigecycline (**Supplemental Figure 1**). In this respect, it should be noted that the 4-ring backbone of KBP-7072 is a structural feature that is common to both the parental Tetracycline (TET) molecule and with the first third generation tetracycline developed, Tigecycline (TIG) (**Figure 1**). Accordingly, this 4-ring moiety of KBP-7072 makes similar interactions with the binding pocket as described in previous work on Tetracycline or Tigecycline (8, 9, 12). Briefly, as illustrated in **Figure 3A-B** and summarized schematically in **Supplemental Figure 1**, these shared interactions include (i) the coordination of a Mg2+ ion between the polar groups of rings B and C, (ii) the coordination of a second Mg2+ ion by the phosphate backbone of G966 in helix 31 and ring A, (iii) hydrogen bonds to residues G1198 and C1195, and (iv) a stacking interaction between C1054 and ring D. In addition to these common interactions, the strong density observed in both the initial Fo-Fc difference map (**Figure 2B**) and final 2F_o_-F_c_ electron density map (**Figure 2C**) allowed us to confidently model and describe interactions for the unique 3-methyl-3-azabicyclo[3.1.0]hexane sidechain moiety that differentiates KBP-7072 from other Tetracycline derivatives. Specifically, the tertiary amine in the 3-methyl-3-azabicyclo[3.1.0]hexane sidechain is clearly positioned in the complex to form a hydrogen bond with the 2’ hydroxyl group of C1054 (**Figure 3A-B,** dotted yellow line) and an ionic hydrogen bond with the phosphate group of G1053 (**Figure 3A-B,** dotted cyan line). Importantly, the potential interactions formed by the sidechain of KBP-7072 (this study) and Tigecycline on the 30S subunit (8) are distinct (**Supplemental Figure 1**). In the case of KBP-7072, these interactions lead the sidechain to assume a bent or kinked conformation on the 30S ribosomal subunit which is different than the extended conformation adopted by the sidechain of Tigecycline in complex with the 30S ribosomal subunit (**Figure 4**; compare orange and green molecules). This bent conformation can be rationalized by the strong ionic hydrogen bond between the tertiary amine in the sidechain of KBP-7072 and the negatively charged 16S rRNA phosphate group of G1053 (**Figure 3**, dotted cyan line) which leads to the bending of the sidechain to favor the positioning of the tertiary amine at a distance of c.a. 3 Å from the phosphate group. This strong electrostatic interaction also allows one to rationalize the observation of clear density for the KBP-7072 sidechain, both in the initial Fo-Fc difference map (**Figure 2B**) and final 2F_o_-F_c_ electron density map (**Figure 2C**) further explaining why, in comparison, the side chain of TIG, which performs with the ribosome a much weaker set of interactions (**Supplementary Figure 1**), is observed in the corresponding electron density maps only at a much lower threshold (see for example **Figure 2A-B** in reference (8)). It should be noted that a bent but distinct conformation has also been observed in the structure of Tigecycline bound to the 70S ribosome (**Figure 4;** purple molecule). In these structures, however, the secondary amine in the *tert*-butylglycylamide sidechain of Tigecycline is not positioned to allow significant direct interactions with the 16S rRNA phosphate backbone and is not discussed in previous Tigecycline-70S structures (9, 12). This difference in the ability of KBP-7072 and Tigecycline to form direct interactions with the 16S rRNA phosphate backbone could stem from the fact that the *tert*-butylglycylamide side chain of tigecycline harbors a peptide bond (**Figure 1**), that should adopt a planar conformation due to its partial double bond character (in conjugation with the heteroaromatic D ring of the drug) and restrict its rotational freedom. Namely, although the side chain of Tigecycline may adopt a bent conformation under certain conditions, this conformation would be hampered by the planar nature of the peptide bond present in the TIG C9 extension while it is freely accessible in KBP-7072 and indeed it predominates in the 30S·KBP-7072 complex, favoring a well-defined, stable and direct interaction with G1053.

**Figure 3:**
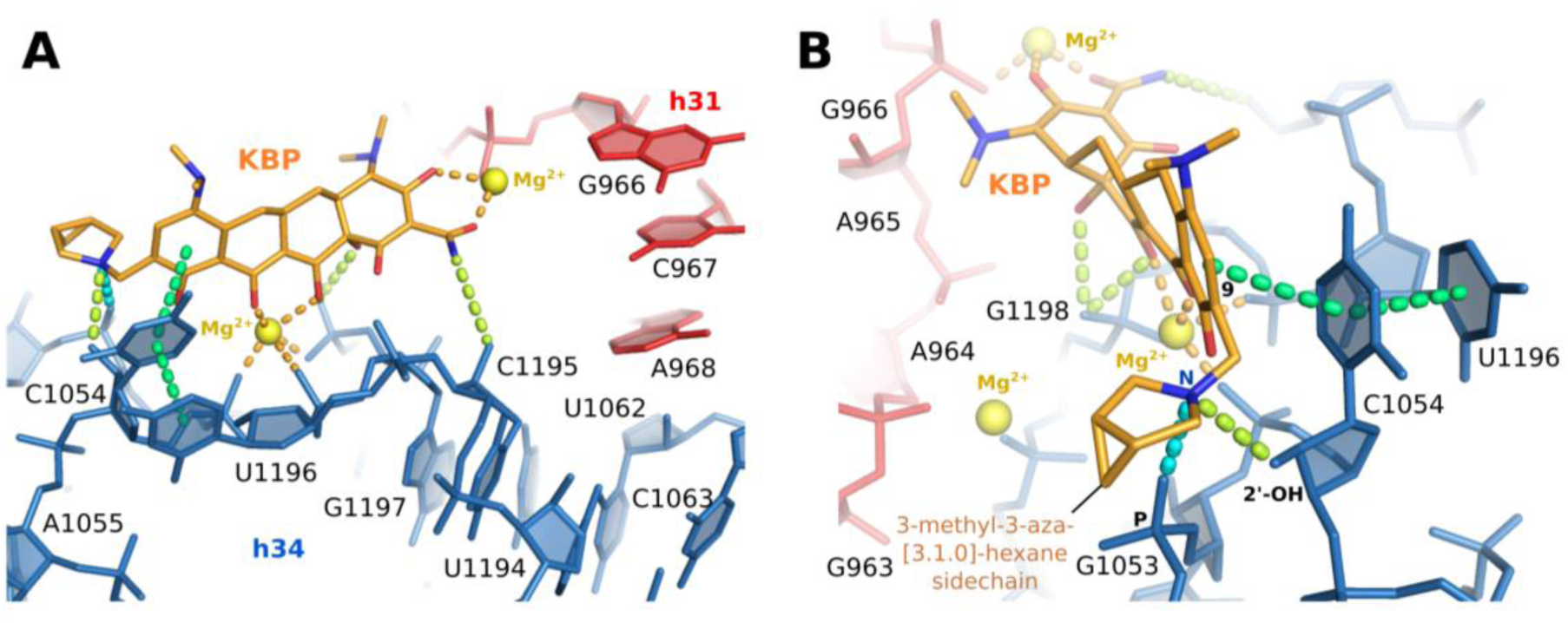
The interactions within the KBP-7072 binding pocket are illustrated on the atomic model in two different orientations (**A-B**). In panels **A-B**, KBP-7072 and the 16S rRNA are colored as described in **Figure 2** with distance-based interactions indicated by dotted lines and specifically hydrogen bonds colored yellow, interactions involving Mg2+ coordination colored orange, stacking interactions colored green and ionic interactions colored cyan. The orientation shown in **Panel B** highlights the hydrogen bond (yellow dotted line) between the tertiary amine (N, blue) present in the 3-methyl-3-azabicyclo[3.1.0]hexane sidechain and the 2’ hydroxyl moiety (2’OH) of C1054, as well as, the ionic hydrogen bond interaction (cyan dotted line) to the phosphate group (P) of G1053. These interactions are in agreement with a protonated state of the tertiary amine, as expected at physiological pH (26) and under the conditions used to form the 30S·KBP-7072 complex (pH 7.6, see **Material and Methods**). The interactions are summarized and compared to the ones observed for Tigecycline when bound to the 30S in **Supplementary Figure 1**.

**Figure 4:**
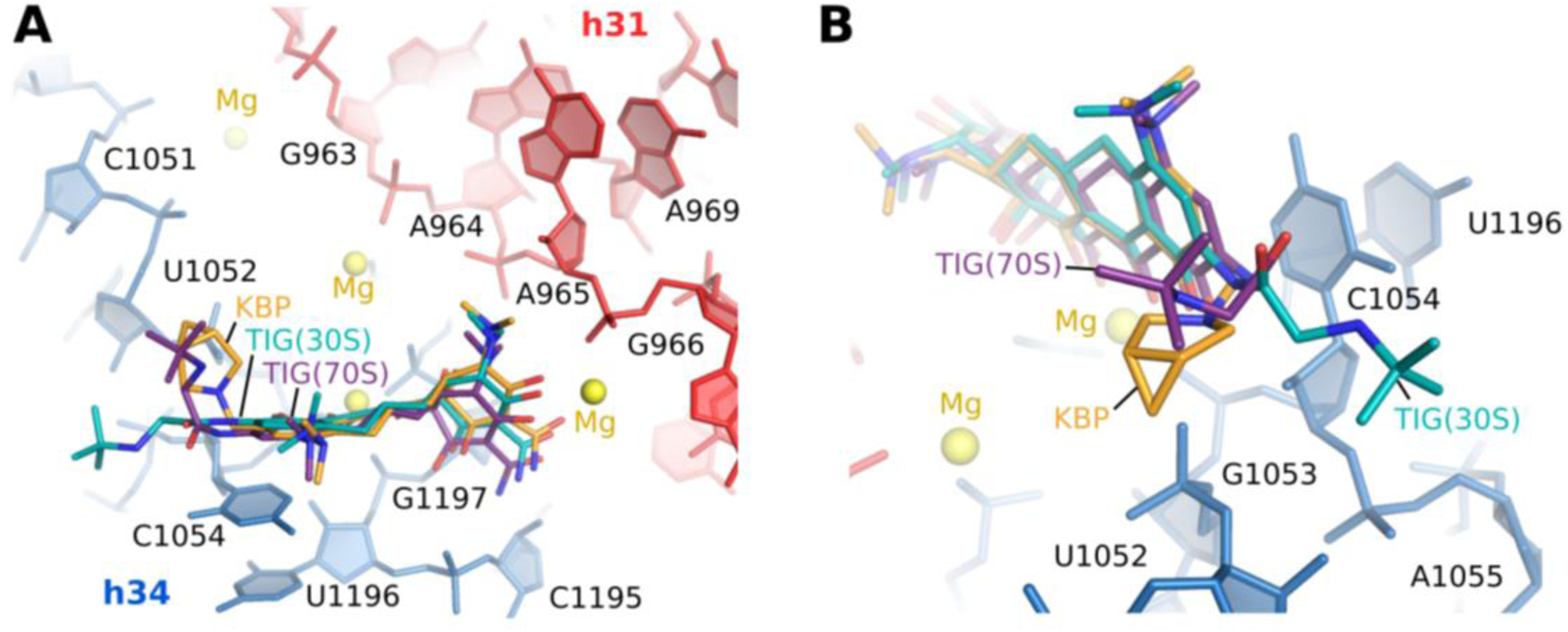
Comparison of the individual binding modes of KBP-7072 (orange) and Tigecycline in complex with the 30S subunit (TIG(30S), green; (8) or in complex with the 70S ribosome (TIG(70S), purple; (9)). Whilst the sidechain of TIG in complex with the 30S (green) shows an extended conformation due to presence of an intact planar peptide bond (being in conjugation to the aromatic backbone ring D, see **Figure 1**) the sidechain of KBP-7072 (orange), bearing two rotatable bonds, around the sidechain methylene group (see **Figure 1**), is instead found in a bent conformation projecting upwards to G1053 in the 30S structure. TIG in the 70S complex (purple) assumes a similar but not identical bent conformation (compare orange and purple molecules in panel B) where, most importantly, the sidechain of TIG is not predicted to form a direct interaction with G1053 (9, 12) which could potentially influence the strength of the interaction between the sidechain and binding pocket.

## DISCUSSION

The X-ray crystal structure of the 30S-KBP-7072 complex reveals that KBP-7072 binding is characterized by a core set of interactions common to all tetracyclines and a unique set of interactions formed by the C9 side chain that distinguishes the modern tetracycline derivatives with C9 extensions. The set of core interactions are responsible for KBP-7072 binding to the ribosomal A-site in a position termed the primary tetracycline binding pocket (**Figure 2A**). Binding to this active site is consistent with KBP-7072’s mode of action being defined as a protein synthesis inhibitor like the typical tetracyclines. Namely, as seen in **Figure 5,** when bound in this position KBP-7072 would overlap with the incoming amino-acyl tRNA (A-tRNA) and sterically hinder the A-tRNA from accessing the decoding site thus blocking protein synthesis. Moreover, as in the case of Tigecycline´s C9 extension (**Figure 5C**), the bulky 3-methyl-3-azabicyclo[3.1.0]hexane sidechain of KBP-7072 (**Figure 5B)** would further contribute to the overlap with the aminoacyl-tRNA potentially enhancing the inhibitory activity of KBP-7072. As seen in **Figure 6,** the bulky side chain of KBP-7072 (**Figure 6B)** could also play a role in overcoming some forms of tetracycline resistance, comparable to that of Tigecycline´s C9 extension (**Figure 6C)**, which restricts the access of ribosomal protection proteins (RPP), like Tet(M), to the primary tetracycline binding site (13). Finally, the distinct chemical nature of the C9 extension and additional interactions formed between this extension and the 16S rRNA (**Figures 3** and **Supplementary Figure 1**) provide a structural basis to differentiate the activity of KBP-7072 from other tetracycline derivatives. For example, the ability of KBP-7072 to form a strong electrostatic interaction with G1053 distinguishes it from Tigecycline where this interaction is not observed in either the 30S or 70S complex structures (8, 9, 12). In this respect, the specific set of interactions observed for KBP-7072 expands the interaction potential of the primary tetracycline binding pocket providing a novel structural framework for the preparation of even more potent tetracycline derivatives.

**Figure 5:**
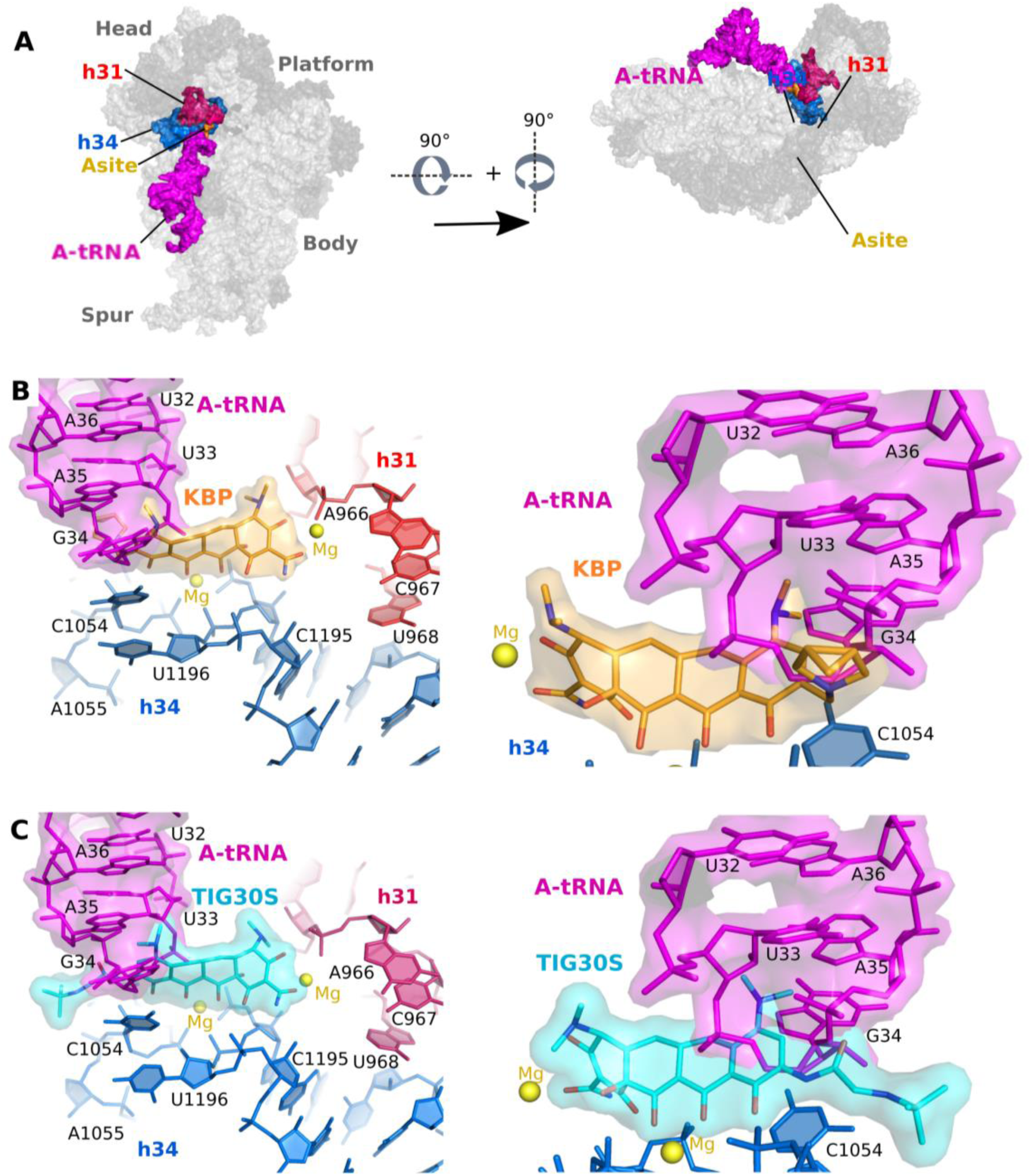
The positions of the primary Tetracycline/KBP-7072 and aminoacyl-tRNA (A-tRNA, magenta) binding site are shown on the structure of the 30S subunit (**A**) with 16S rRNA elements forming the Tetracycline/KBP-7072 binding pocket, namely helix 31(h31) and helix 34 (h34) colored red and blue, respectively and important residues for A-tRNA binding colored yellow. (**B** and **C**) A structural alignment of the A-tRNA (magenta; PDB ID: 5UYL) onto the 30S-KBP-7072 complex highlights the steric clash between the A-tRNA and the antibiotic (KBP-7072, panel **B**; Tigecycline-30S, panel **C**) that allows the drug to inhibit A-site tRNA binding and block protein synthesis. In panels **B** and **C** (left) the structure is shown with the 30S subunit in the orientation seen in panel **A** (right). In the right of these panels the view is from the reverse orientation and highlights the clash between the sidechain and A-tRNA.

**Figure 6.**
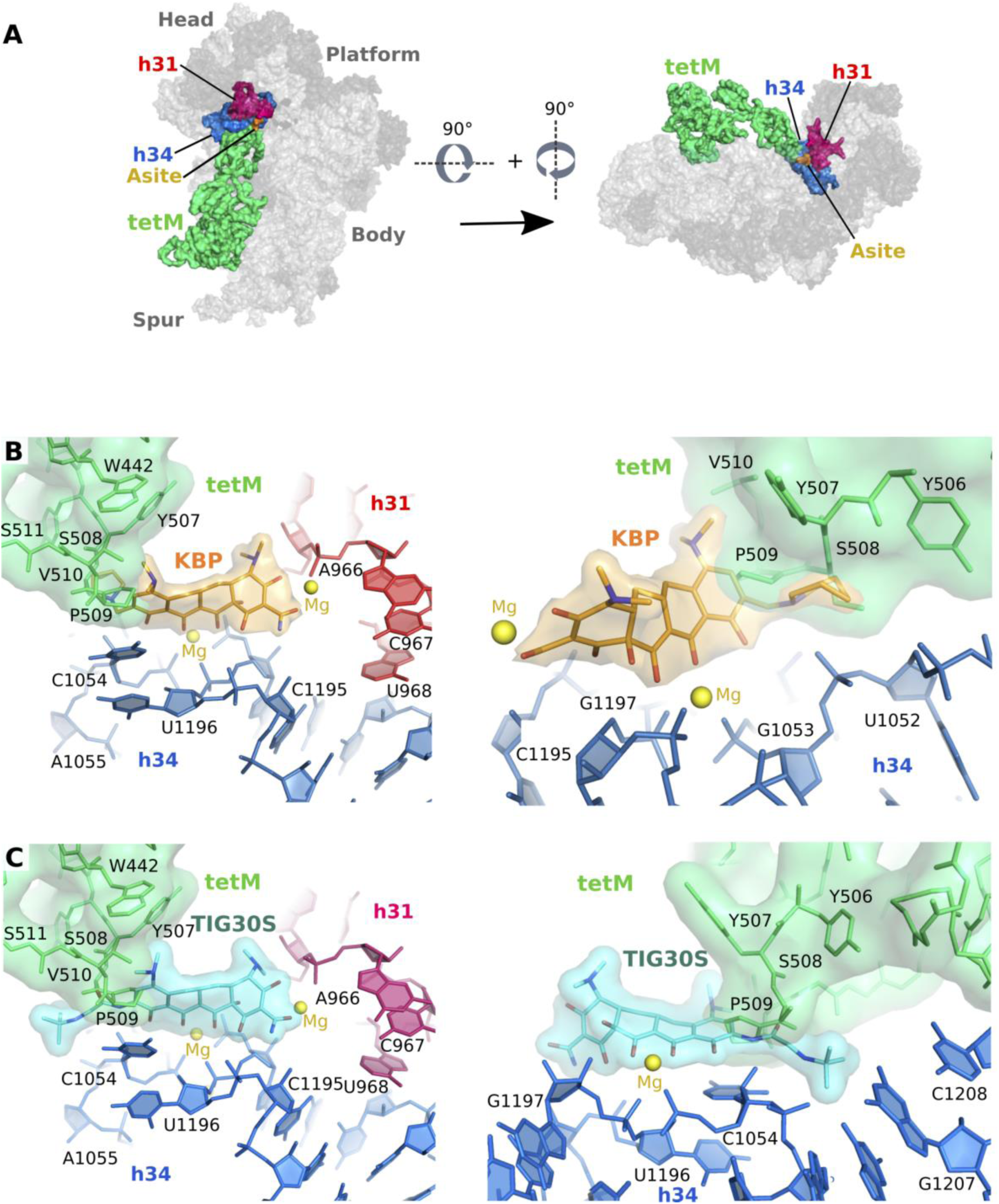
A structural alignment of the ribosome bound Tet(M) structure (PDB ID: 3J9Y) with the 30S-KBP-7072 (**B**) and 30S-Tigecycline (**C**) structures. 16S rRNA elements forming the KBP-7072 binding pocket, namely helix 31 (h31) and helix 34 (h34) are colored red and blue, respectively, and residues corresponding to Tet(M) are colored lime green. This structural alignment highlights the steric clash between Tet(M) and the antibiotic KBP-7072 (panel **B)** and Tigecycline (panel **C**). In panels **B** and **C** (left) the structure is shown with the 30S subunit in the orientation seen in panel **A** (right). In the right of these panels the view is from the reverse orientation as seen on the left and highlights the clash between the sidechain and Tet(M).

## Acknowledgments

We are particularly grateful to KBP BioSciences Co., Ltd. for providing us with their proprietary compound KBP-7072. This study could not have been performed without the expert assistance of the staff at the XALOC beamline at ALBA (Spain), the i04-1 beamline at Diamond Light Source (UK), and the id29 and id30b beamlines at the European Synchrotron Radiation Facility (France). This work was supported by KBP Biosciences Co., Ltd. and grants from the Ministerio de Economía Y Competitividad Grant (CTQ2017-82222-R and CTQ2014-55907-R to P.F., S.R.C.). SC and PF thank MINECO for the Severo Ochoa Excellence Accreditation (SEV-2016-0644).

---

**Supplementary Figure 1.**
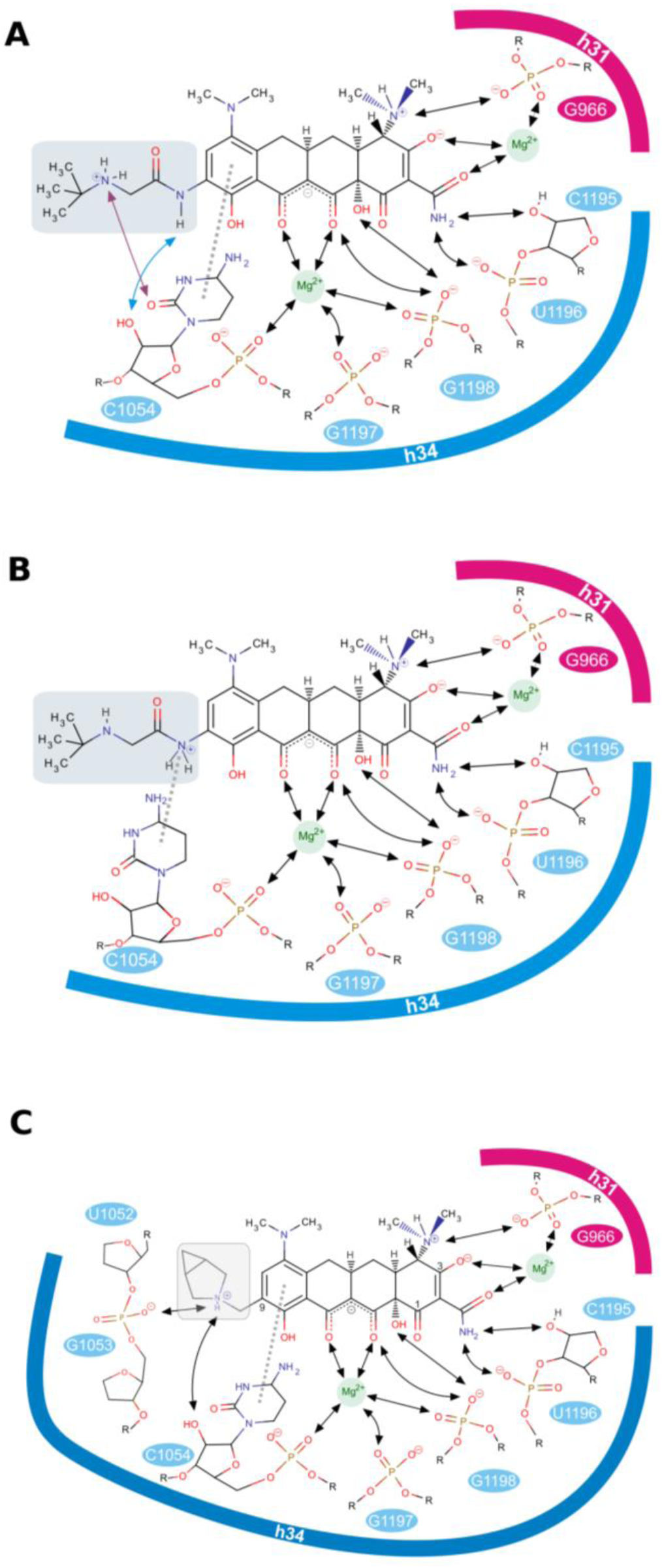
Interactions between the antibiotics and their binding pocket as observed in the structures of Tigecycline bound to the 30S ribosomal subunit (**A;** (8)), Tigecycline bound to the 70S ribosome (**B;** (9)), and KBP-7072 bound to the 30S ribosomal subunit (**C;** this work) are shown schematically. Despite the similarity observed in the three complexes for the interactions of the 4-ring moiety of the drug, the functionally relevant tail of TIG and KBP (highlighted in grey) forms specific sets of interactions in each complex. Notably, only the KBP tail is observed to engage with the ribosome through an ionic hydrogen bond interaction. The strength of this type of interaction (26), allows one to rationalize the well-defined density observed in the KBP-7072 Fo-Fc map (**Figure 2B**).

